# Vertical transmission of tissue microbiota in *Caenorhabditis elegans*

**DOI:** 10.1101/2021.12.06.471348

**Authors:** Jun Zheng, Xin Meng, Jiahao Fan, Dong Yang

**Author notes:** **Correspondence and requests for materials** should be addressed to D.Y. These authors contributed equally: Jun Zheng, Xin Meng.

## Abstract

The past forty-five years has witnessed *Caenorhabditis elegans* as the most significant model animal in life science since its discovery seventy years ago^1,2^, as it introduced principles of gene regulated organ development, and RNA interference into biology^3-5^. Meanwhile, it has become one of the lab animals in gut microbiota studies as these symbionts contribute significantly to many aspects in host biology^6,7^. Meanwhile, the origin of gut microbiota remains debatable in human^8- 11^, and has not been investigated in other model animals. Here we show that the symbiont bacteria in *C. elegans* not only vertically transmit from the parent generation to the next, but also distributes in the worm tissues parallel with its development. We found that bacteria can enter into the embryos of *C. elegans*, a step associated with vitellogenin, and passed to the next generation. These vertically transmitted bacteria share global similarity, and bacterial distribution in worm tissues changes as they grow at different life stages. Antibiotic treatment of worms increased their vulnerability against pathogenic bacteria, and replenishment of tissue microbiota restored their immunity. These results not only offered a molecular basis of vertical transmission of bacteria in *C. elegans*, but also signal a new era for the mixed tissue cell-bacteria multi-species organism study.

---

Gut microbiota plays a vital role in many aspects of host physiology, and becomes more so as research inputs increases^12^. The sterile womb doctrine deduces that human gut microbes are established after delivery^13,14^. However, once the gut microbiota lost its richness due to unbalanced diet or antibiotic treatment, it seems impossible to restore them by supplementing the host with routine nutritional interventions^15^. This raises the questioning that environment could not reseed bacteria into the gut, and ultimately leads to the tracing back of these bacteria in an individual: what is their origin? Efforts have been tried to find evidences for vertical bacterial transmission via human umbilical cord blood, amniotic fluid, or placenta^8-10^. Meanwhile, new evidences point out bacteria in the placenta are merely contaminations in the sample collection process, laboratory reagents, or sequencing instrument^11^. Owing to the bacterial world we are residing in, it seems impossible to prove the sterility of any tissue under current experimental conditions^16^.

*C. elegans* has been used as the animal model for many studies, including genetic regulation of organ development and programmed cell death, RNA interference, and development of green fluorescent proteins^3-5,17^. And it has been recently employed in the gut microbiome research^6,7,18,19^. Microbiota in worms from different microbial environments resembles each other and share a core flora^20^, and the bacterial composition is influenced by their developmental stage and genotype^21^. The diverse worm microbiota exhibits significant heterogenicity within *E. coli* species that affect host longevity via communicating with colonic acid^22^. Growing in the lab for decades on monoxenic cultures (mostly *E. coli* OP50), the N2 wild type worm harbors some bacteria (genus *Exiguobacterium, Mucilaginibacter*, and *Virgibacillus*) that does not exist in the produce enriched, native microbiota restored soil^20^. Thus, a similar question arises that where are the worm gut bacteria from except *E coli*?

Taking advantage of many technical conveniences with nematode *C. elegans* as the model animal, we first tried to examine whether there is a vertical transmission of bacteria in this model as in many other animals^23^. Here we found there is not only core bacteria in the nematode embryo but also in its tissues, and these tissue microbiota plays an important role in host defense against pathologic bacteria. Additionally, the maternal transmission of tissue microbiota is associated with vitellogenin, probably during the embryo formation process.

## Embryo microbiota of *C. elegans*

To verify the maternal transmission of bacteria in C. elegans, one has to prove there are bacteria in the worm embryo as previously verified in *Haemonchus contortus*^24^. The traditional bacteria probe Eub338 was used in this study^25^, and it is examined capable of visualizing both gram-negative and -positive bacteria with different efficiency though (Extended Data Fig.1a-b)^26^. We found there are bacteria inside the *C. elegans* embryo *in situ*, most likely intercellular ones (Fig.1a-b, Supplementary Video 1). Isolated, surface sterilized embryo also identified the presence of bacteria within it (Supplementary Video 2)

**Fig. 1.**
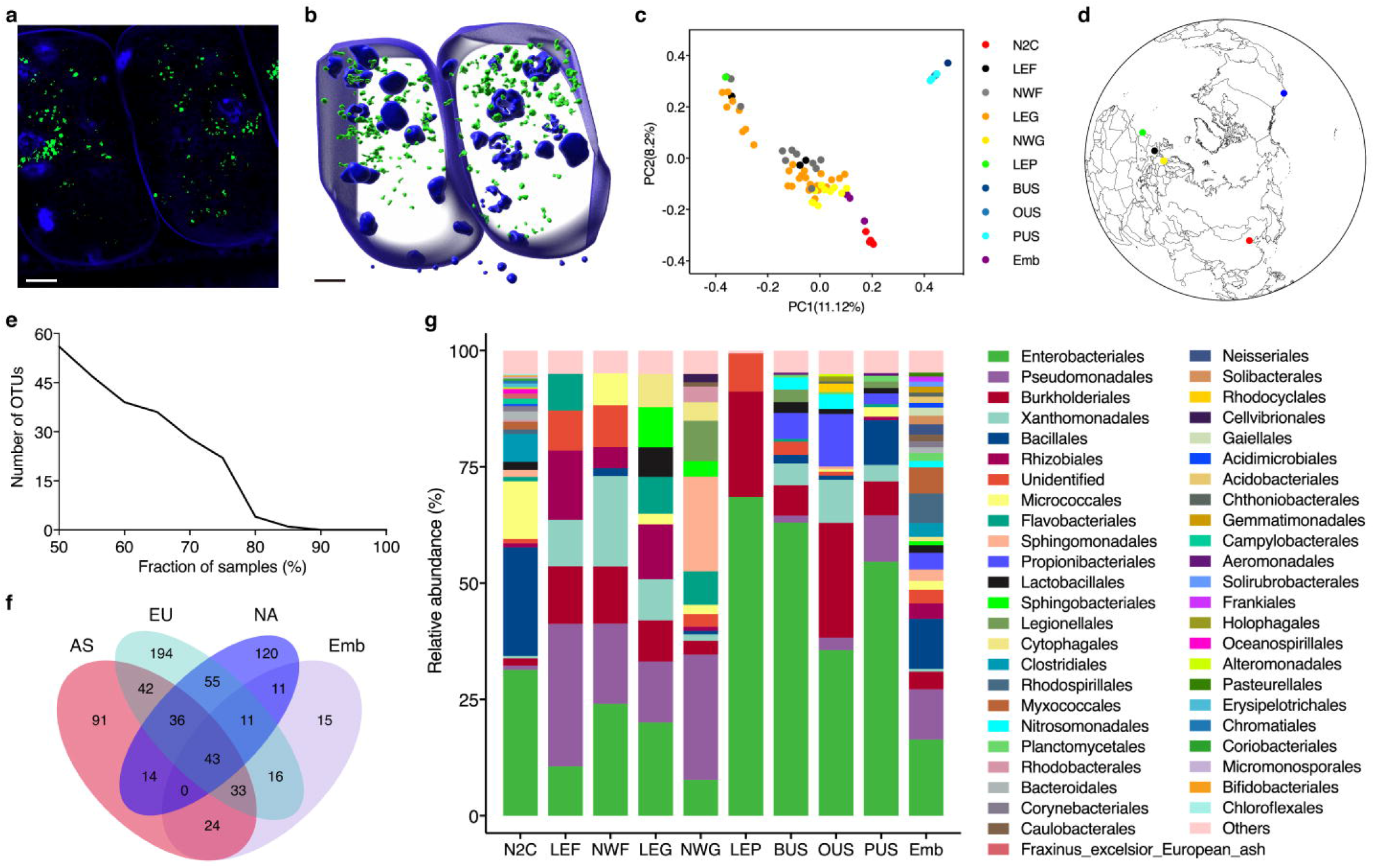
Vertical transmission of bacteria in *C. elegans*. **a**, the confocal microscopic image of worm embryos inside a hermaphrodite, blue is the DAPI stained worm cell and green is the probe labelled bacterial DNA. **b**, volumetric reconstruction of the worm embryos. Scale bars indicate 5μm. **c**, the Bray-Curtis PCoA of worldwide *C. elegans* microbiota, x-axis is 11.12% explained variance and y-axis is 8.2% explained variance. N2C (n=5) is the N2 nematode in Chinese labs. LEF and NWF (n=14) are lab enriched and natural wild worms in France. LEG and NWG (n=40) are lab enriched and natural wild worms in Germany. LEP (n=1) is lab enriched worm in Portugal. BUS, OUS, and PUS (n=9) are banana, orange, and potato fed worms in the U.S.. Emb (n=3) is the worm embryo in Chinese labs. **d**, the locations of worm samples in **c**. Red indicates China, blue indicates U.S., black in indicates France, yellow indicates Germany, and green indicates Portugal. **e**, core OTUs across 50%-100% of all the samples in the above. **f**, Venn diagram of common bacteria in the level of genus of all the samples in the above. **g**, bacterial composition in the level of order, the groups are the same as in **c**.

We sequenced the 16S rRNA gene in the isolated embryo and the N2 laboratory worms in our lab, and compared with these from the worm gut microbiome available to date^20,21^. Principle coordinate analysis indicated the gut microbiota of worms from China, France, Germany, Portugal, and the U.S. mainly clustered according to their geographic origin (Fig. 1c-d). Interestingly, microbiota in the embryo clustered centering among these global worms, indicating a universal similarity of bacteria in the embryo with these in the worms. Although no common OTUs were observed among all worms (Fig.1e), there are 43 common genera between worms from different continent and the embryos (Fig.1f). Microbiota composition analysis indicated that Enterobacteriales, Pseudomonadales, and Burkholderiales are the common orders among all the samples (Fig.1g)

### Tissue microbiota of *C. elegans*

Routine FISH processing enabled us to visualize the gut bacteria localized along the intestine (Extended Data Fig.2a-b, Supplementary Video 3), with the same pattern as the ingested, red fluorescent protein labelled *E. coli* (Extended Data Fig.2c-d, Supplementary Video 4-6). Interestingly, there are also bacteria detected at the oocyte and spermatheca (Extended Data Fig.3, Extended Data, Supplementary Video 1,7). To distinguish the gut bacteria and these in worm tissues, much effort was devoted to optimize the FISH procedure (Additional Discussion 1). We found that bacteria in the worm gut could be removed by washing (Extended Data Fig.4), and FISH procedure on worms with their tails cut off improves visualization of bacteria in worm tissues while the wound did not allow foreign bacteria entering into the worm tissue (Extended Data Fig.5).

Sex dependent FISH procedure enables us to clearly localize tissue bacteria distribution. For hermaphrodites, the probe was incubated with worms at 80°C then 37°C, followed by washing at 37°C (see methods for detail). This enabled us to visualize bacteria not only in the embryos but also in different tissues (Extended Data Fig.6). In the gut microbiota removed hermaphrodites, bacteria were seen in the pharynx, body wall muscle, and embryos (Fig. 2a, c, Extended Data Fig.7). However, this procedure only permits visualization of bacteria in the male worm spermatheca (Extended Data Fig.8). For male worms, optimized FISH procedure was incubating the probe with worms at 46°C followed by washing at 48°C. This allowed us not only localize bacteria in the spermatheca but also in the pharynx and body wall muscle (Fig. 2b, d, Extended Data Fig.9). However, hermaphrodites performed with the later FISH setup barely detects bacteria in the embryos (Extended Data Fig.10).

**Fig. 2.**
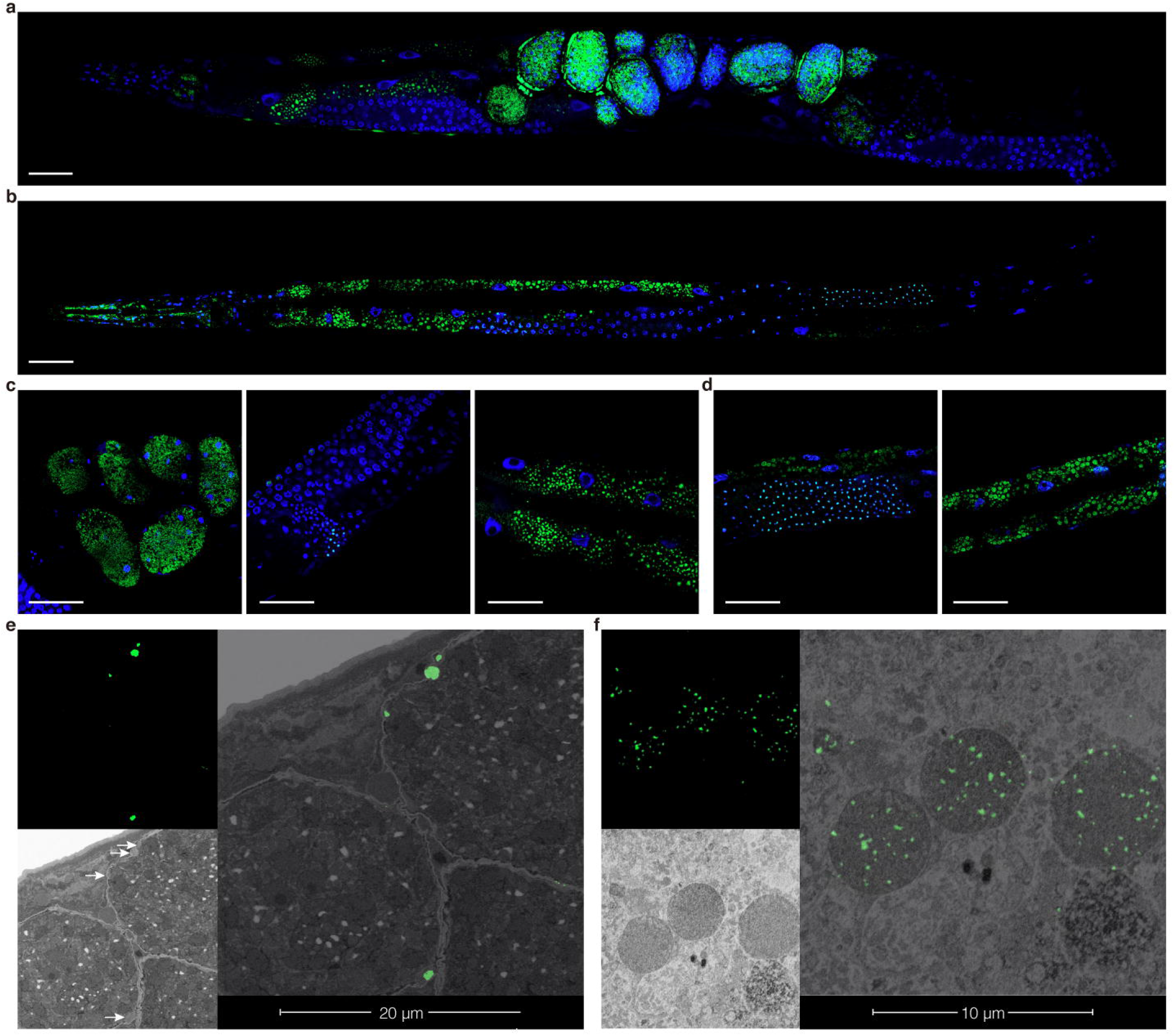
Tissue bacteria in *C. elegans*. **a**, FISH of tissue bacteria in a hermaphrodite under confocal microscope, blue is the DAPI stained worm cell and green is the probe labelled bacteria DNA. **b**, FISH of tissue bacteria in a male under confocal microscope. **c**, FISH of organs in a hermaphrodite, from left to right are embryos, spermatheca, dorsal and ventral body wall muscle. **d**, FISH of organs in a male worm, from left to right are the gonad, dorsal and ventral body wall muscle. Color in **b**-**d** are the same as in **a**. Scale bars indicate 30μm. **e**, CLEM of tissue microbiota in a hermaphrodite embryo, upper left is the FISH image of bacteria in the embryos, lower left is the SEM image of the same site and whit arrow indicates the corresponding position of the fluorescence signal. Right is the overlay of light and electron microscopy, and green is the anti-digoxin antibody labelled bacteria. **f**, CLEM of tissue microbiota in a male worm yolk, upper left is the FISH of bacteria in the yolk, lower left is the SEM of the same site, and right is the overlay. Green indicates the same as in **e**.

To further localize bacteria in worm tissues, correlative light and electron microscopy (CLEM) was applied to detect tissue bacteria in *C. elegans*. Briefly, digoxin labelled nucleic acid probe was used to hybridize *in situ* with tissue bacteria in the gut microbiota free, tail cut worms. These worms were then subjected to ultrathin sectioning, labelled with fluorescent anti-digoxin antibody, and observed sequentially with confocal light microscopy and electron microscopy (Extended Data Fig.11). Proof-of-concept experiment was performed with both gram-negative and -positive bacteria (Extended Data Fig.12a-c), and this procedure unequivocally prevents contamination of foreign bacteria into the worm tissue during experimental operation (Additional Discussion 2). CLEM helped us identified bacteria inside the embryo (Fig.2e, Extended Data Fig.12d) in hermaphrodites and inside the sperm cells in male worm gonad and yolk (Fig.2f, Extended Data Fig.12e).

We then tried to visualize the tissue microbiota distribution dynamics during *C. elegans* development. Bacteria seemed occupying all over the worm body at the L1 stage (Extended Data Fig.13a-b, Extended Data Fig.14a-b, Supplementary Video 8), and the tissue microbiota became less populated at the L2 stage (Extended Data Fig.13c-d). The tissue bacteria continued to vanish at the L3 (Extended Data Fig.13e) and L4 stages (Extended Data Fig.13f), with a few left in the lateral tissues (Extended Data Fig.14c-d, Supplementary Video 9). Our images did not allow us for more precise description of the tissue microbiota distribution.

### Tissue microbiota assisted worm immunology

To examine the physiological significance of tissue microbiota, we isolated the *C. elegans* embryos and soaked them with different antibiotics before they entered into the L1 stage, by which the gut microbiota is not involved in this intervention. Worm growth under the pathological *Enterococcus faecalis* challenging did not yield significant differences between those treated with saline and ampicillin or chloramphenicol, while these treated with kanamycin exhibited significantly shortened lifespan (Log-rank test, *P*<0.05, Extended Data Fig.15). The more challenging pathological *Pseudomonas aeruginosa* was applied^27^, and it was found that worms treated with kanamycin and chloramphenicol both exhibited significant shortened life span compared with these treated with saline and ampicillin (Log-rank test, *P*<0.05, Fig.3a). Worm tissue microbiota were sequenced, and PCoA indicated that tissue microbiota of saline treated worms almost overlapped with ampicillin treated one while kanamycin and chloramphenicol treated ones deviated from others (Fig.3b).

**Fig. 3.**
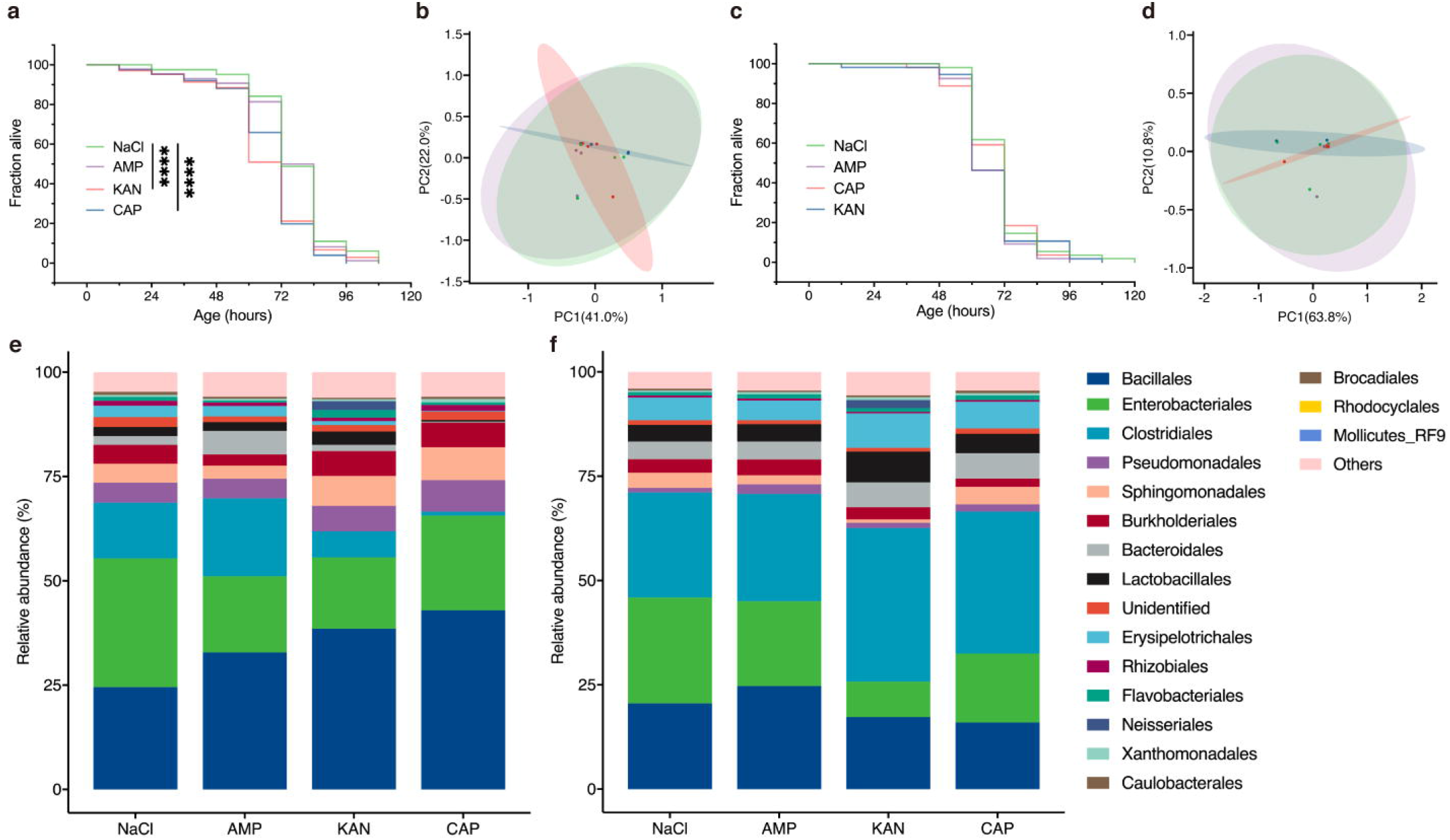
Tissue bacteria related immunity in *C. elegans*. **a**, life span of worms treated with different antibiotics including ampicillin, kanamycin, chloramphenicol with saline as the control, and under the challenge of *Pseudomonas aeruginosa*. Worms treated with kanamycin and chloramphenicol exhibited significantly shortened life span compared with the control group (Log-rank test, *p*<0.0001). **b**, Bray-Curtis PCoA of the tissue microbiota in different antibiotic treated *C. elegans*, x-axis is 41.0% explained variance and y-axis is 22.0% explained variance. Green indicates saline, purple indicates ampicillin, red indicates kanamycin, and blue indicates chloramphenicol treated worms. **c**-**d**, corresponding analysis of different antibiotic treated worms, then replenished with tissue bacteria. There is no significant difference among life spans of different antibiotic treated worms in **c. e-f**, the top 15 tissue bacterial composition at the order level of the antibiotic treated worms, and the antibiotic treated, then tissue bacteria replenished worms, respectively. The groups are the same as in **a**.

To further prove the role tissue microbiota played in *C. elegans* immune system, we replenished bacteria from worms to these treated with different antibiotics. It was found that the life span differences between worms treated with kanamycin and chloramphenicol, and these treated with saline and ampicillin disappeared (Fig.3c). PCoA indicated that the tissue microbiota of antibiotic treated, then bacteria replenished worms clustered closer to the saline treated ones (Fig.3d). Bacteria analysis indicated that the relative abundance of orders Bacillales, Sphingomonadales, and Burkholderiales increased after kanamycin treatment and decreased after following tissue bacteria replenishment. While those of orders Clostridiales, Erysipelotrichales, Bacteroidales, Brocadiales, and Rhodocyclales decreased after kanamycin treatment and increased after bacteria replenishment. Meanwhile, the relative abundance of orders Bacillales and Burkholderiales increased after chloramphenicol treatment and decreased after following bacteria replenishment. While those of orders Clostridiales, Erysipelotrichales, Bacteroidales, Brocadiales, and Lactobacillales decreased after chloramphenicol treatment and increased after following bacteria replenishment (Fig.3e-f). It is worthy to mention that the antibiotic treatment did not permanently alter the worm tissue microbiota as they restored after couple of generations (Extended Data Fig.16)

### *Vit-2* associated tissue microbiota vertical transmission

We often observed the cooccurrence of bacteria with the yolk in the embryo, thus it was speculated that yolk component might be involved in the vertical transmission of tissue microbiota. It was reported that vitellogenin in fish binds to bacteria as a multivalent pattern recognition receptor^28^, thus we turned to RNA interference on the yolk protein yp170B, the protomer of the homodimeric yolk lipoprotein complex to examine its role in bacterial vertical transmission^29^. For the F0 generation on which the RNAi was applied, there is no significant differences between life spans of worms supplied with *E. coli* OP50, *E. coli* HT115, *E. coli* HT115 harboring empty vector, or *E. coli* HT115 harboring RNAi plasmid under pathogenic *Pseudomonas aeruginosa* challenging (Fig.4a). PCoA of tissue microbiota indicated the almost overlapped clustering between RNAi treated and control group (Fig.4b), and composition analysis showed that there was no significant difference between species at the order level (Fig.4c).

**Fig. 4.**
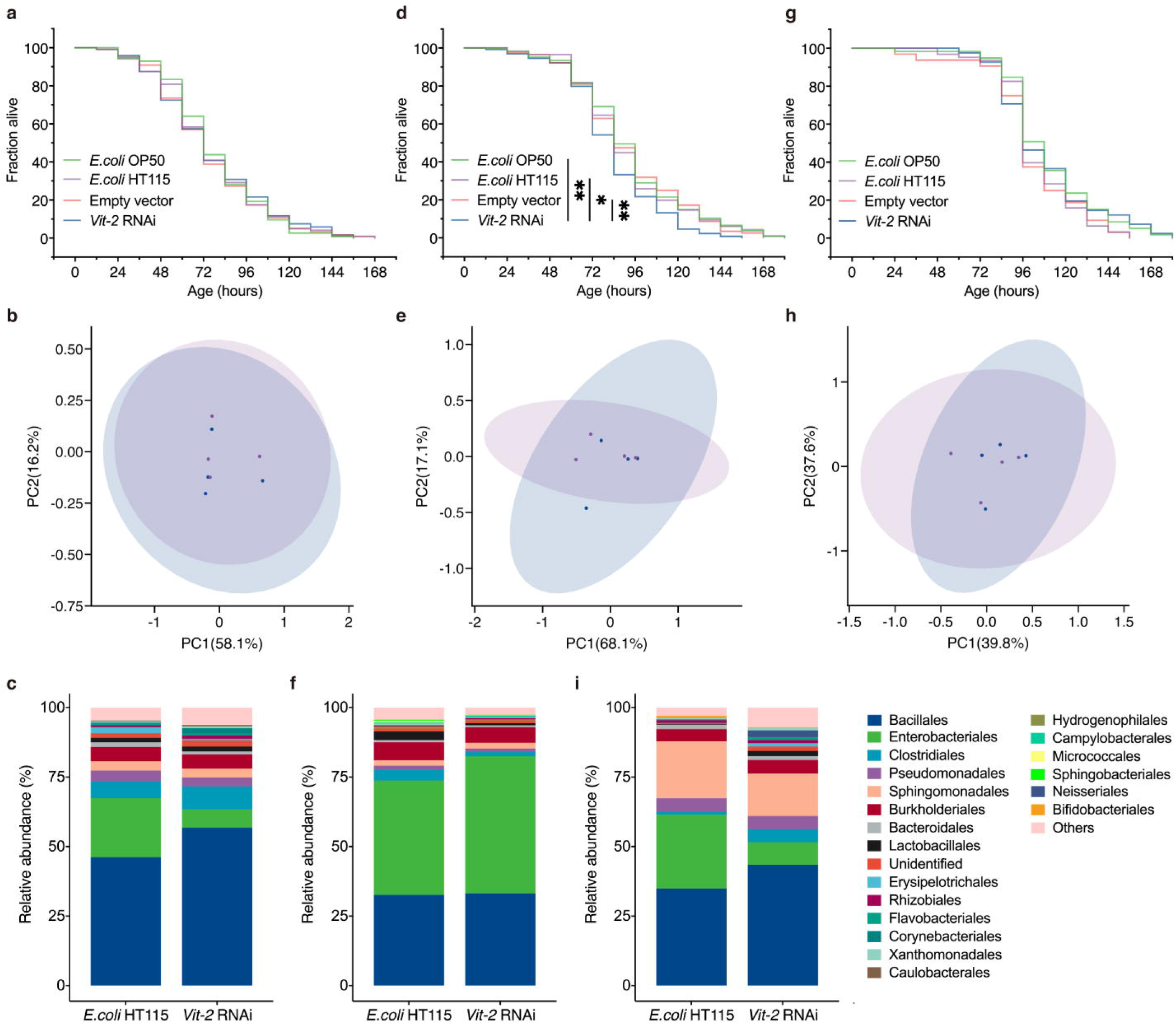
Vitellogenin mediated vertical microbiota transmission in *C. elegans*. **a**, life span of RNAi worms under the challenge of *Pseudomonas aeruginosa*. Green, purple, red, and blue indicate worms were fed with *E. coli* OP50, *E. coli* HT115, *E. coli* HT115 harboring empty vector, and *E. coli* HT115 harboring *Vit*-2 RNAi plasmid. There is no statistical significance among the life spans of differently treated worms **b**, Bray-Curtis PCoA of the tissue microbiota in different RNAi treated *C. elegans*, x-axis is 58.1% explained variance and y-axis is 16.2% explained variance. Purple indicates worms fed with *E. coli* HT115 and blue indicate worms fed with *E. coli* HT115 harboring *Vit*-2 RNAi plasmids. **c**, top 15 bacterial composition at the order level, the groups are the same as in **b. d**-**f** same analysis of the progeny of worms treated with RNAi. Progeny of worms fed with *E. coli* HT115 harboring *Vit*-2 RNAi plasmids exhibited significantly shortened life span compared with the other groups (Log-rank test, *indicates *p*<0.05 and ** indicates *p*<0.01). **g**-**i**, same analysis of progeny of worms treated with RNAi then replenished with tissue bacteria. There is no statistical significance among the life spans of differently treated worms in **g**.

However, in the progeny F1 generation under the same pathogen challenging, RNAi treated *C. elegans* exhibited significant shortened life span compared with control worms (Fig.4d). PCoA indicated tissue microbiota clustering of these two groups deviated from each other as compared with the F0 generation (Fig. 4e). Among orders with most intensive relative abundance changes, relative abundance of orders Enterobacteriales, Sphingomonadales increased in the RNAi treated worms compared with these supplied with *E. coli* HT115 while these of orders Clostridiales, Lactobacillales, Burkholderiales, Erysipelotrichales decreased (Fig. 4f). Again, we replenished tissue bacteria to a parallel batch of F1 worms, and found that their life span discrepancy under pathogenic challenging disappeared (Fig.4g). Their tissue bacterial composition is to certain extent restored to the F0 generation thus microbiota clustering overlapped closer (Fig. 4h). Relative abundance of Enterobacteriales, Sphingomonadales decreased in the RNAi worms replenished with tissue bacteria while these of orders Clostridiales, Lactobacillales, Burkholderiales, Erysipelotrichales increased (Fig. 4i).

## Discussion

Bacteria were detected in the eggs, the intestinal epithelia and oocytes of nematodes *Xiphinema brevicollum* and *Xiphinema americanum*^30^, in the gut, uterus, tissue, and inside the eggs of adult nematode *Haemonchus contortus*^24^, and in the ovary of nematode *Acrobeles sp*.^31^. Our results showed that in the model animal *C. elegans*, there is bacteria in the embryo with shared core species to be passed to their progeny. Specific interest into detection of tissue bacteria rose when we observed detection of bacteria in the spermatheca and embryos with EUB338, and in the ovary as reported^31^. As the obligate intracellular bacteria *neorickettsia spp*. reported to be maintained within all life cycle stages of another laboratory model *Plagiorchis elegans*^32^, and here we also observed intracellular EUB338 signal in *C. elegans* in its various tissues. Vitellogenin is synthesized in the intestine and transported into maturing oocytes by endocytosis^33^, and it is also capable of binding bacteria^28^. Our results indicated vitellogenin is associated with bacterial vertical transmission, as the worm immuno-compromise brought by RNAi treatment on vitellogenin only occurs in the progeny generation and could be restored by microbiota replenishment. Vit-2 protein distributes all over the worm tissues as yolk accumulates^34^, and so are the bacteria it binds as we observed.

Subgroups of *Pseudomonas fluorescens* inhibits pathogen growth^35^, *Bacillus megaterium* enhanced worm immunology via decreasing worm reproduction, and *Pseudomonas mendocina* did so associated with *pmk-1*-dependent immune gene expression^36^. Although these above bacteria help host against pathogens in different mechanisms, no evidence shows they are functioning in the worm gut cavity. Despite vitellogenin assisted *C. elegans* immunology against pathological bacteria via the steroid-signal pathway^37^, here we find vertical transmission and tissue distribution of *Pseudomonas fluorescens*, another subgroup of which exhibit immunology depends on host-mediated mechanisms^35^. It is highly likely this subgroup is functioning in the worm tissue via communication with *C. elegans* cells.

Differential vitellogenin provisioning is responsible for phenotypic variation in the isogenic *C. elegans* whilst the mechanism remains unknown^38^. Here we identified that vitellogenin is associated with vertical transmission of tissue bacterial to the progeny which could profoundly impact animal immunity, and possibly other physiology. Traditionally viewed as an autonomous process directed by the genome solely, animal development now appeals a reexamination in the presence of hologenomes as a holobionts^16,39,40^.

## Methods

### *C. elegans* and bacteria

The wild-type N2 *C. elegans* were maintained on standard nematode growth medium (NGM) seeded with *Escherichia coli* OP50 as previously described, except the temperature was set at 20°C^2^. Both the worm and the OP50 cells were kindly provided by Prof. Guangshuo Ou at Tsinghua University. *E. Coli* BL21 constitutively expressing red fluorescence protein (RFP) was kindly provided by Prof. Yanling Hao at China Agricultural University in which sequence coding red fluorescence protein was inserted between the restriction nuclease site *Nco*I and *BamH*I of a pET9d plasmid. *Enterococcus faecalis* ATCC 29212 was purchased from China General Microbiological Culture Collection Center. *Pseudomonas aeruginosa* PA14 was kindly provided by Dr. Kun Zhu in the Institute of Microbiology, Chinese Academy of Sciences. *E. coli* HT115 strain was purchased from Shanghai Weidi Biotechnology Co., Ltd (Shanghai, China).

### Gut microbiota positioning

RFP labeled *E. Coli* was inoculated in the LB medium and cultured overnight. On a 60 mm NGM dish, 50μL of the above *E. Coli* was seeded at 37°C for 5-8 hours. Around 100 *C. elegans* adult worms were picked and maintained on the above lawn for one day, and then hermaphrodites and male worms were separately picked, washed with M9 buffer once and 1×PBST buffer three times. Worms were then heat-shocked at 60°C for 1 min before collected by centrifugation. For 10μL of the worm sample, add 5μL anti-fading mounting medium with 5μL 4’, 6-diamindino-2-phenylindole (DAPI, Solarbio Life Sciences, Beijing China), transfer to a slide and cover with a glass coverslip. To prevent evaporation of the mounting solution, the edges of the cover glass was sealed with nail polish (OPI Products Inc., N. Hollywood, CA, USA).

### Removal of gut bacteria by washing

Worms were collected by flushing with M9 buffer, washed with 1×PBST buffer three times, and heat-shocked at 60°C for 1 min. Worms were then collected by centrifugation, incubated in 1mL 4% PFA for 45min on a rocking shaker, washed three times with 1×PBST buffer, resuspended in 1mL 1×PBST buffer, and washed on a rocking shaker at for 0h, 1h, or 2h, respectively. The worms were collected by centrifugation, washed again with fresh 1×PBST buffer, and subjected to DNA probe *in situ* hybridization as described in the following.

### Nucleic acid probe in situ hybridization

In this study, two types of nucleic acid probes were used. Fluorescent probe for direct confocal microscopic observation, Eub338 (5’ - GCTGCCTCCCGTAGGAGT-3’)^26^ with Alexa Fluor 488 dye covalently linked to its 5’ end, which detects almost all bacteria, was synthesized by Sangon Biotech Co., Ltd. (Shanghai, China). The other probe used in the following CLEM experiment was Eub338 with digoxin covalently attached to its 5’ end (Sangon Biotech, Shanghai, China).

Routine hybridization procedure is as the following. Hermaphrodites and male worms were separately picked, washed with 1×PBST buffer three times, and heat-shocked at 60°C for 1 min. Worms were collected by centrifugation, incubated in 1mL 4% PFA for 45min on a rocking shaker, washed three times with 1×PBST buffer, resuspended in 100μL hybridization buffer (900mM NaCl, 20mM Tris-Cl pH7.2, 0.01% SDS, 20% formamide), and transferred to a 200μL PCR tube pre-rinsed with hybridization buffer containing 0.5% Tween-20. The DNA probe was added to a final concentration of 1μM, and the animals were stained at 46°C for 1.5h on a rocking shaker for male, or 80°C in a water bath for 30min and then 37°C on a rocking shaker for 1.5h for hermaphrodites. Worms were collected by centrifugation and washed on a rocking shaker with cleaning solution (180mM NaCl, 20mM Tris-Cl pH7.2, 0.01% SDS) at 48°C for 30min for male, and 37°C for 30min for hermaphrodites. Worms were washed again with the cleaning solution, and collected by centrifugation. For 10μL sample, add 5μL of anti-fading mounting medium with 5μL DAPI, transfer to a slide and cover with a glass coverslip. To prevent evaporation of the mounting solution, the edges of the cover glass was sealed with nail polish (OPI Products Inc., N. Hollywood, CA, USA). Worms at different development stage, isolated worm embryos, or bacteria (Bacteria *E. coli* OP50 and *E. faecalis* ATCC 29212) were following the FISH procedure for male.

For tissue microbiota FISH experiment, the tail of each worm was cut off with a scalpel under a compact stereo microscope (Olympus SZ61, Japan). Worms were then collected and washed with 1×PBST buffer as inspired by a previous study^41^. These steps were performed after the PFA fixation and PBST cleaning, but before the hybridization buffer addition.

The DNA probe hybridized samples were subjected to confocal microscopic imaging (with DAPI) or embedding according to the DNA probe used.

### Foreign bacterial contamination test

The RFP labelled *E. coli* was cultured, collected by centrifugation, washed three times with 1×PBST buffer, and used to substitute the DNA probes in the above *in situ* hybridization procedure for male. After the cleaning step, anti-fading mounting medium with DAPI were added and the sample were covered with a glass coverslip for confocal microscopic observation.

### Ultrathin section preparation

In the high-pressure freeze (HPF) procedure, samples were carefully loaded into a type A specimen carrier (200-μm well) containing 10% BSA in M9 buffer, and covered with a top hat with a thin coating of 1-hexadecane. The two carriers were hold firmly together, and loaded into the sample holder for HPF (Leica HPM100, Germany). Following HPF, the fast-frozen samples were immersed into a freezing tube containing 1% osmium tetroxide in 100% acetone, and placed into the freeze substitution (FS) device (Leica EM AFS, Germany) with the following parameters: T1 = -90°C for 60 h, S1 = 3°C/h, T2 = -60°C for 10 h, S2 = 3°C/h, T3 = -30°C for 14 h, then slowly warmed to 4°C (4°C/h). Following FS, samples were rinsed three times in 100% acetone, 15 min each at 4°C, and once again at room temperature. Samples were then transferred into a new 2ml Eppendorf tube and infiltrated in a graded mixture (1:5, 1:3,1:1, 3:1) of resin (SPI, Resin Mixture: 16.2 ml SPI-PON812, 10 ml DDSA, 10 ml NMA,1.5%BDMA) and acetone mixture, then changed to 100% resin four times for 2 days on rotator. Finally, samples were embedded and polymerized 12h at 45°C, and 48h at 60°C. The ultrathin sections (200nm thick) were sectioned with microtome (Leica EM UC7) on a glass slide.

### Ultrathin section labelling

Under a microscope, the ultrathin section was moved to a glass slide with 200μL 1mg/mL BSA solution with the Perfect loop (70940, Head Biotechnology Co., Ltd., Beijing, China) and incubated for 30min at room temperature. Then the section was moved to another glass slides with 100μL, 1μM anti-digoxin antibody in 1×PBS buffer and incubated at 37°C for 3h. The section was then moved to another glass slide with 200μL, 1mg/mL BSA and incubated for 10min, before finally to a glass slide with 200μL H2O for 10min. The stain and washed section was then moved to a cover slide with 50μL H2O, air-dried at room temperature, and subjected to confocal microscopic and electron microscopic observation.

### Confocal microscopy and imaging analysis

Alexa Fluor 488, DAPI, and RFP signal distributions in the worm were captured with an inverted Olympus FV1000 confocal microscope system with 10× and 60× objectives, or Zeiss LSM980 confocal microscope system with 40×, 63× and 100× 1.40-NA objectives. The excitation wavelength was set at 488/488nm, 405/405nm, and 635/594nm with FV1000/LSM980 for Alex Fluor488, DAPI, and RFP signal, respectively. For 3D visualization, acquisition was performed in 0.5 or 1 μM z-steps depending on the thickness of the specimen, and the Imaris software package (Bitplane AG, Zurich, Switzerland) was used to visualize z-stack, reconstruction of the 3D architecture of the samples and video production.

### Scanning electron microscopy

Ultrathin sections were double stained with uranyl acetate and lead citrate, coated with carbon in a high vacuum evaporator (Denton Vacuum 502B, NJ, USA), and examined on a scanning electron microscope (FEI Helios Nanolab 600i, Oregon, USA) in the immersion high magnification mode with a CBS detector at 2kV and 0.6nA. For CLEM, the light microscopy images and electron microscopy images of the same area were carefully aligned.

### Antibiotic treatment

On an NGM plate with majority of adult hermaphrodites harboring eggs, the worms were collected by flushing M9 buffer to the plates, transferred to a sterile centrifuge tube, and washed three times with M9 buffer. The residual suspension of worms in 100μL M9 buffer was added sequentially with 600μL the same buffer, 100μL 5M NaOH solution, and 200μL NaClO solution (10% available chlorine), inverted vigorously until all worms lysed, and centrifuged at 1300×g for 1 min. The embryos were washed with M9 buffer until pH neutral and resuspended in 400μL buffer.

M9 buffer of 3 mL was added to pre-rinsed 60 mm petri dishes, and 6μL of each 100g/mL saline, ampicillin, kanamycin, and chloramphenicol (Shanghai Macklin Biochemical Co., Ltd., Shanghai, China) solutions was added to the solution for a final concentration of 200μg/mL, respectively. The embryos in different solutions were incubated at 20°C for hatching. The solutions were then transferred to centrifuge tubes after 24 hours, centrifuged at 1300×g for 2 min, and the supernatant was discarded. Worms at L1 stage were washed three times with M9 buffer, resuspended in M9 buffer and transferred to NGM plates, and referred as the F0 generation. The F0 generation was removed from the plates by gently flushing the plates with M9 medium after they laid the embryos, and the offspring was referred as the F1 generation and transferred to new NGM plates. Similarly, the offspring of the F1 generation was referred as the F2 generation. F0 worms at L4/young adult stage were subjected to life span assay as described in the following. F0, F1, and F2 worms were subject to tissue microbiota analysis.

### RNA interference

The pUC57 plasmid harboring sequences targeting the *vit2* gene was synthesized in Tsingke Biological Technology, Beijing, China (Supplementary information), transformed into the DH5α *E. coli* cell (TransGen Biotech, Beijing, China) and amplified, extracted and digested with *Hind* III and *Xho* I restriction enzymes (Takara Biomedical Technology, Co., Ltd., Beijing, China). The L4440 linear vector (MiaoLing Plasmid Sharing Platform, Wuhan, China) was digested with the same restriction enzymes, mixed with the *vit2* sequences in equal molar ratio, and mixed with the T4 ligase (Vazyme Biotech Co. Ltd., Nanjing, China) before incubation at 16°C for 1h. The ligation product was transformed into the DH5α cell and single colonies were sequenced at Tsingke Biological Technology, Beijing, China. Cells harboring correct plasmid were cultured and pL4440-*vit2* was extracted with the TIANprep Mini Plasmid Kit (Tiangen Biotech Co. Ltd., Beijing, China). Meanwhile, the empty pL4440 plasmid was prepared. The pL4440 and pL4440-*vit2* plasmids were heat-shock transferred into the HT115 (DE3) competent cells (Shanghai Weidi Biotechnology Co., Ltd., Shanghai, China), respectively. HT115 cells harboring pL4440 or pL4440-vit2, were all cultured in LB medium containing 50μg/mL ampicillin, and HT115 cells harboring no plasmids were also plated, then cultured on LB medium. The RNAi plates were made by seeding these cultures on the NGM plates containing 4mM isopropyl-β-d-thiogalactopyranoside (IPTG, Shanghai Macklin Biochemical Co., Ltd., Shanghai, China), culturing at 37°C for 48h.

Worm embryos were obtained by the above bleaching method, washed to pH neutral, added to each RNAi plate, incubated at 20°C and referred as the F0 generation. The F0 worms were gently removed from the plates by flushing with M9 medium, and the left embryos were collected, washed, transferred to NGM plates for hatching, and referred as the F1 generation. F0 or F1 worms at L4/young adult stage were subjected to life span assay as described in the following, and tissue microbiota analysis.

### Somatic bacteria replenishment

Worms at L4 stage were collected by flushing them from the NGM plates with M9 buffer, centrifuged to remove the supernatant buffer, washed three times with the M9 buffer, grinded with a sterilized grinding rod (rinsed with 10% hydrochloric acid overnight, washed to pH neutral, and pasteurized at 121°C for 20min). The grinding-lysed worms examined with no intact animals were spread over the NGM plates (covered with OP50 *E. coli*). Replenishment was performed by transferring the antibiotic treated worm embryos (the F0 generation), or the embryos of the RNAi treated worms (the F1 generation) to the above NGM plates containing tissue bacteria and cultured at 20°C till the worms developed to the L4/young adult stage for the following lifespan assay, and tissue microbiota analysis.

### Lifespan assays

Worm life span assays were performed at 20°C. Synchronization of worm populations was performed by placing L4/young adult worms (treated with antibiotics, RNAi, or the F1 generation of RNAi treated worms) on NGM plates seeded with *Enterococcus faecalis* ATCC 29212 or *Pseudomonas aeruginosa* PA14. Worms were changed to new plates every day for elimination of confounding progeny and were marked as alive or dead. Worms were scored every 12/24 hours, counted as dead if they did not respond to repeated prods to with a platinum pick, and censored if they crawled off the plate or died from vulval bursting. In each life span assay, 50-60 worms were allowed to grow in a plate, and the assays were repeated for triplicates. The data were plotted into the Kaplan-Meier Survival curves with GraphPad Prism, and the statistical significance was determined by Mantel-Cox Log-rank.

### Microbiota DNA isolation and 16S rRNA sequencing

Worms or isolated embryos were washed 3 times with M9 buffer before subjected to whole microbiome extraction with the Rapid Soil DNA Isolation Kit (Sangon Biotech, Shanghai, China) according to manufacturer’ s instruction. Worms were washed 3 times with M9 buffer, incubated with 4% PFA on a rocking shaker for 45min, then washed again with water for 3 times, incubated in water on a rocking shaker for 2 hours, and finally washed with water once before tissue microbiome extraction with the same DNA isolation kit. Amplification of the V3-V4 region of the 16S rRNAs of the microbiota were performed with the barcoded primers 336F (5’ -GTACTCCTACGGGAGGCAGCA-3’) and 806R (5’ - GTGGACTACHVGGGTWTCTAAT-3’). The PCR reaction contains a total of 20μL volume 1×FastPfu buffer, 10ng template DNA, 250μM dNTP, 0.1μM each above primer, and 1 U FastPfu DNA polymerase (TransGen Biotech, Beijing, China). PCR was performed at 95°C for 2 min, followed by 30 cycles of 95°C for 30s, annealing at 55°C for 30s, 72°C for 30s and a final extension at 72°C for 5 min. The amplicons were detected with 2% agarose gel electrophoresis and purified with AxyPrep DNA Purification kit (Axygen Biosciences, Union City, CA, USA). Purified amplicons were pooled in equal molar and paired-end sequenced (2×300) on an Illumina MiSeq platform (Illumina Inc., San Diego, CA, United States) at Allwegene Technology Inc. (Beijing, China).

### Microbiota analysis

The sequence files from Berg et. al. was downloaded from http://metagenomics.anl.gov/, referred to Orange enriched worms from US (OUS1-3), Potato enriched worms from US (PUS1-3), and Banana enriched worms from US (BUS1-3) as originally O1w, O2w, O3w, P1w, P2w, P3w, B1w, B2w, and B3w^20^. The sequence files from Dirksen et. al. was downloaded from the European Nucleotide Archive, and named as Lab Enriched worms from France (LEF1-3), Lab Enriched worms from Germany (LEG1-30), Lab Enriched worms from Portugal (LEP1), Natural worms from France (NWF1-11), Natural worms from Germany (NWG1-10) by the order of location, year and month of collection^21^. Consistent with these above-mentioned samples, microbiome including the tissue and gut ones from our lab is referred as N2 worms from China (N2C), and microbiome in embryos isolated from our lab is referred as Emb. For the Pair-End data, the 300bp reads were truncated at any site that obtained an average quality score of <20 over a 50bp sliding window, and the truncated reads shorter than 100bp were discarded with Trimmomatic (v. 0.36)^42^. The truncated data were then merged with a minimum overlap of 10bp and an error matching rate of 0.1 using FLASH (v. 1.20)^43^. Chimeric sequences were identified and removed with UCHIME, and clean tags were clustered into operational taxonomic units (OTUs) with 97% similarity cutoff with VSEARCH (v. 2.7.1)^44^ using the clustering method UPARSE^45^. OTUs were analyzed with QIIME (v. 1.8.0)^46^ for rarefaction analysis and calculation of diversity indices including Principal Coordinates Analysis (PCoA). Taxonomic classification of the representative sequence of each OTU was performed with RDP Classifier^47^ and the Silva (release 128)^48^ 16S rRNA database using confidence threshold of 90%.

## Acknowledgments

This work was supported by the National Laboratory of Biomacromolecules, Institute of Biophysics, Chinese Academy of Sciences (2018kf09). We thank Prof. Guangshuo Ou in Tsinghua University, Prof. Yanling Hao in China Agricultural University, and Dr. Kun Zhu in the Institute of Microbiology, Chinese Academy of Science for offering *C. elegans* worms and bacteria strains, Ms. Yun Feng, Xixia Li, Xueke Tan, Li Wang, and Yan Teng in the Center for Biological Imaging (CBI), Institute of Biophysics, Chinese Academy of Sciences for their help in electron and microscopic imaging, Prof. Hong Zhang and Ms. Huichao Deng in Institute of Biophysics, Chinese Academy of Sciences for discussion. The authors are grateful to Prof. Chih-chen Wang in Institute of Biophysics, Chinese Academy of Sciences for her support and encouragement in our research.

## Author contributions

D.Y. conceived and designed the project. J.Z., X.M., and J.F. carried out the study, overseen by D.Y. and both analyzed the data. D.Y. wrote the manuscript. All authors read and approved the manuscript.

