## Extended Figures for "Vertical transmission of tissue microbiota in *Caenorhabditis elegans*"

### Extended Data Figure Legends

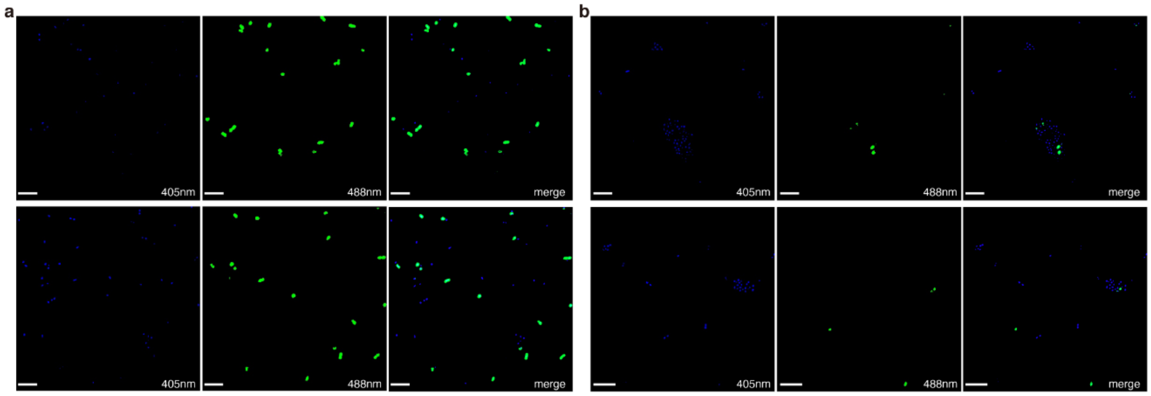


**Extended Data Fig. 1 Fluorescent probe labelled bacteria *in vitro*. a**, The DAPI and Eub338 probe with an Alexa Fluor 488 dye attached were used in the labelling of gram-negative *E. coli* OP50 in solution. Bacteria were stained at 46℃ and washed at 48℃. **b**, the same was used in the labelling of gram-positive *E. faecalis* ATCC 29212 in solution. The left columns are the DAPI signal of stained bacteria with excitation wavelength of 405nm, the middle columns are the Alexa Fluor 488 signal of stained bacteria with excitation wavelength of 488nm, and the right columns are the merge of corresponding left and middle images. Two lines are independent replicates and the scale bars represent 5μm.

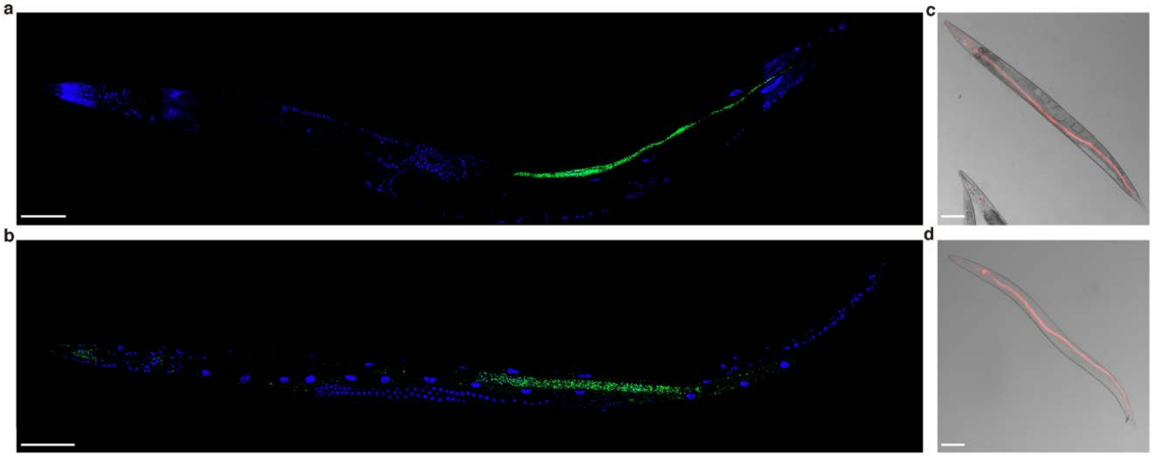


**Extended Data Fig. 2 Positioning of gut microbiota in *C. elegans*. a,** the DAPI and Eub338 probe with an Alexa Fluor 488 dye attached were used to stain an intact hermaphrodite. Worms were stained at 46℃ and washed at 48℃ before observed under the excitation of 405nm and 488nm with a 40× oil immersion objective. **b**, same staining of a male worm as in **a**. Scale bars represent 50μm. **c**, confocal fluorescent imaging of a hermaphrodite fed with red fluorescent protein labelled *E. coli*, and the worms were observed under the excitation wavelength of 635nm with a 10× objective. **d**, the same of a male worm as in **c**. Scale bars represent 100μm.

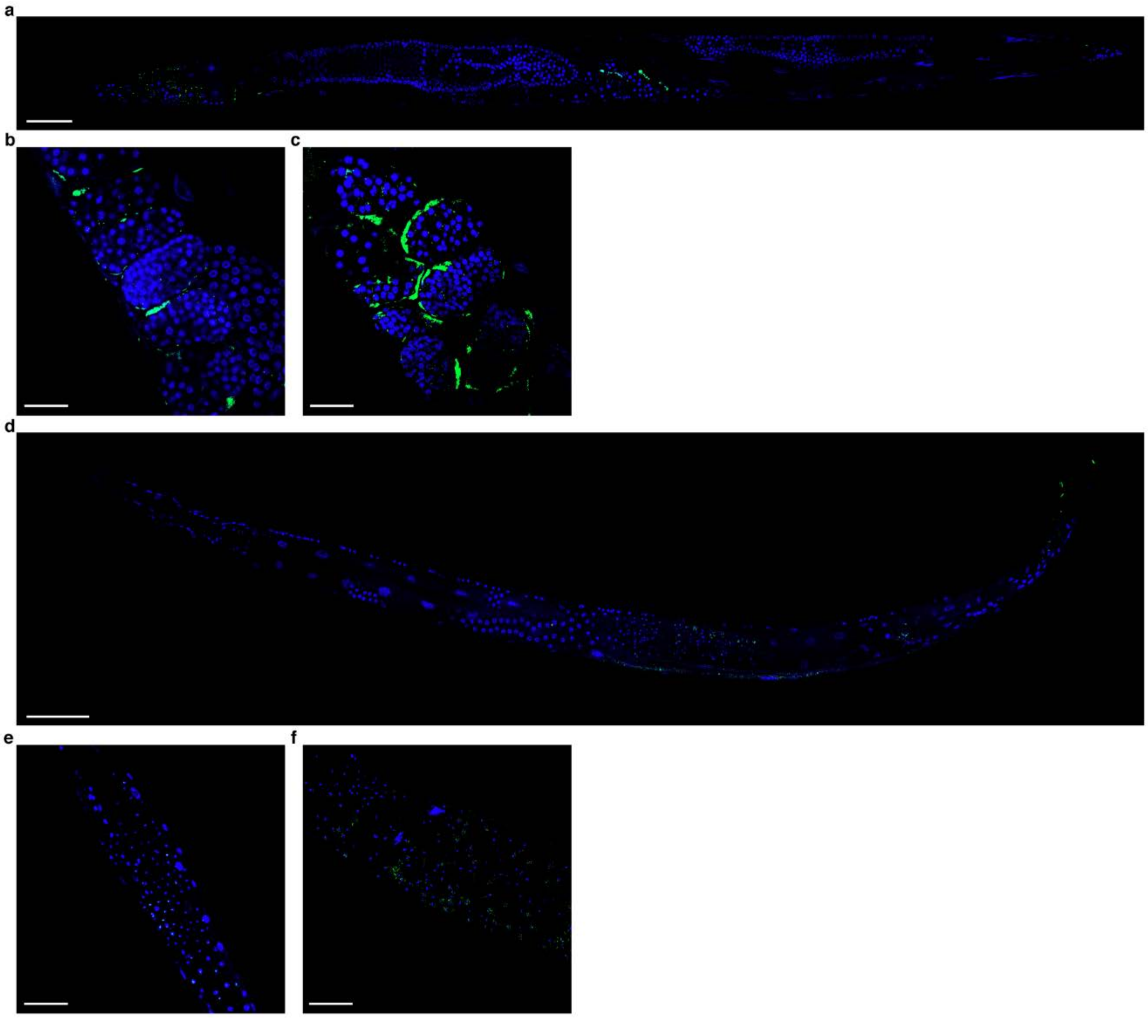

**Extended Data Fig. 3 FISH of tissue bacteria in intact *C. elegans*. a**, visualization of tissue bacteria in a hermaphrodite stained with DAPI and Alexa Fluor 488 dye labelled Eub338. The worms were stained at 46°C and washed at 48°C. Blue is the DAPI signal and green is the signal of Eub338. The image was obtained under the excitation of 405nm and 488nm with a 40× oil immersion objective. Scale bar represents 50μm. **b-c**, oocytes in hermaphrodites observed with the same procedure as in **a**, and scale bars represent 20μm. **d**, visualization of tissue bacteria in a male worm with the same procedure as in **a**, and scale bar represents 50μm. **e-f**, spermatheca in male worms observed with the same procedure as in a, and scale bar represents 20μm.

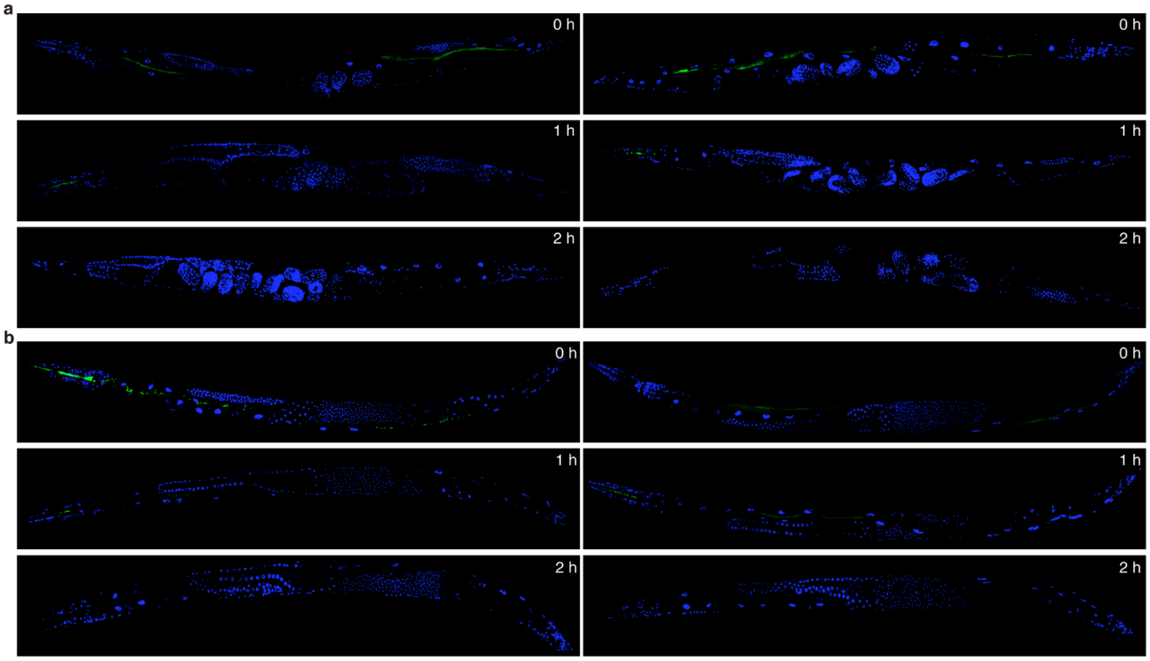


**Extended Data Fig. 4 Washing off of gut microbiota in *C. elegans*. a**, the gut bacteria imaging by staining a hermaphrodite with Alexa Fluor 488 dye labelled Eub338 and DAPI with a 10× objective of warms before washed, washed for 1h and 2h, respectively. **b**, the same washing experiment with a male worm. Different columns are independent replicates.

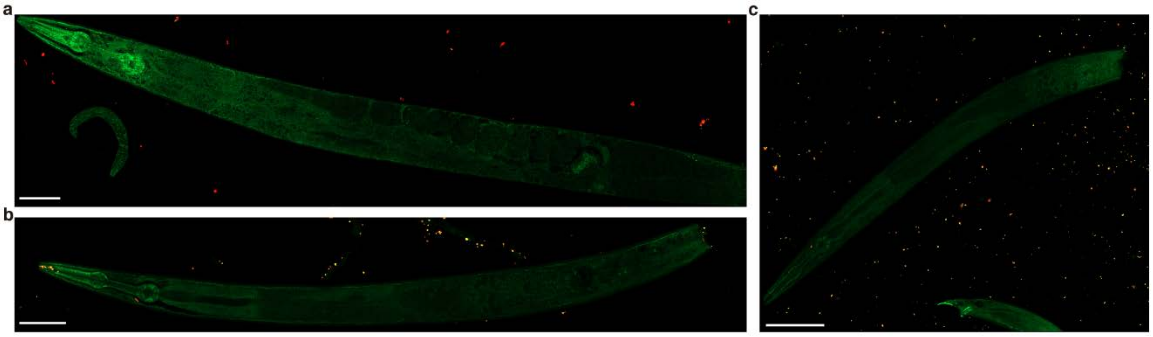

**Extended Data Fig. 5 No contamination of foreign bacteria into tail-cut *C. elegans*. a**, image of a hermaphrodite, whose tail was cut off, with red fluorescent protein labelled *E. coli* used as probes to stain following the FISH procedure as heat-shocked at 80°C, stained at 37°C, and washed at 37°C. The excitation wavelength of the confocal fluorescent microscope was set at 488nm and 594nm, and image was obtained with a 40× oil immersion objective. Scale bar represents 50μm. **b-c**, image of a male worm, whose tail was cut off, with red fluorescent protein labelled *E. coli* used as probes to stain following the FISH procedure as stained at 46°C and washed at 48°C. The confocal microscopic set up was the same as in **a**, and scale bars represent 50μm.

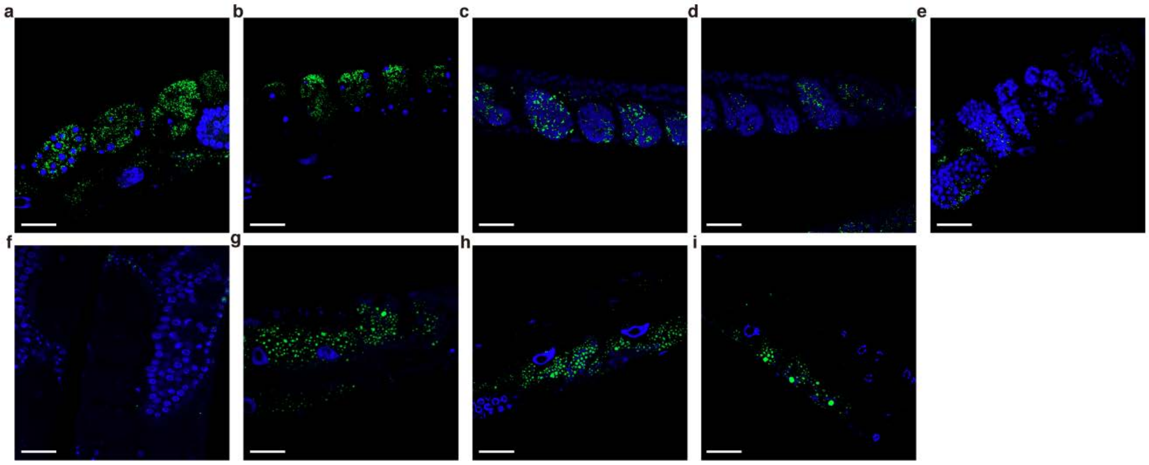


**Extended Data Fig. 6 Visualization of tissue bacteria in the organs of a hermaphrodite.** The tissue bacteria imaging in **a-e** the embryos, **f** the spermatheca, and **g-i** the body wall muscle by staining a hermaphrodite, whose tail was cut off, with DAPI and Alexa Fluor 488 dye labelled Eub338. Worm was heat-shocked at 80℃, stained at 37℃, and washed at 37℃. The excitation wavelength of the confocal fluorescent microscope was set at 405nm and 488nm, and image was obtained with a 40× oil immersion objective Scale bars represent 20μm.

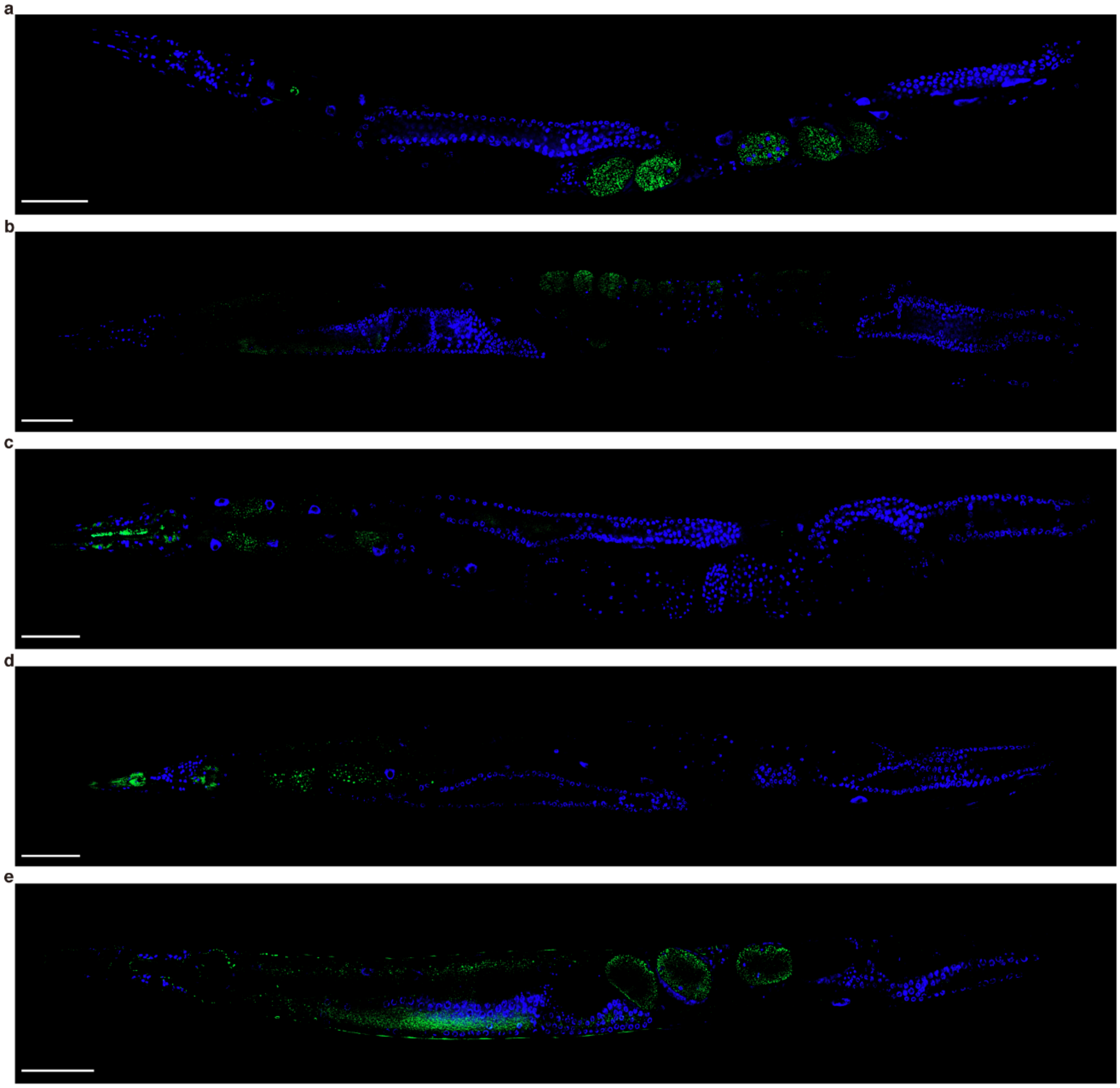


**Extended Data Fig. 7 Visualization of tissue bacteria of a hermaphrodite. a-e**, five typical images of the tissue bacteria imaging by staining a hermaphrodite, whose tail was cut off, with DAPI and Alexa Fluor 488 dye labelled Eub338. Worms were heat-shocked at 80°C, stained at 37°C and washed at 37°C. The excitation wavelength of the confocal fluorescent microscope was set at 405nm and 488nm, and image was obtained with a 40× oil immersion objective. Scale bars represent 50μm.

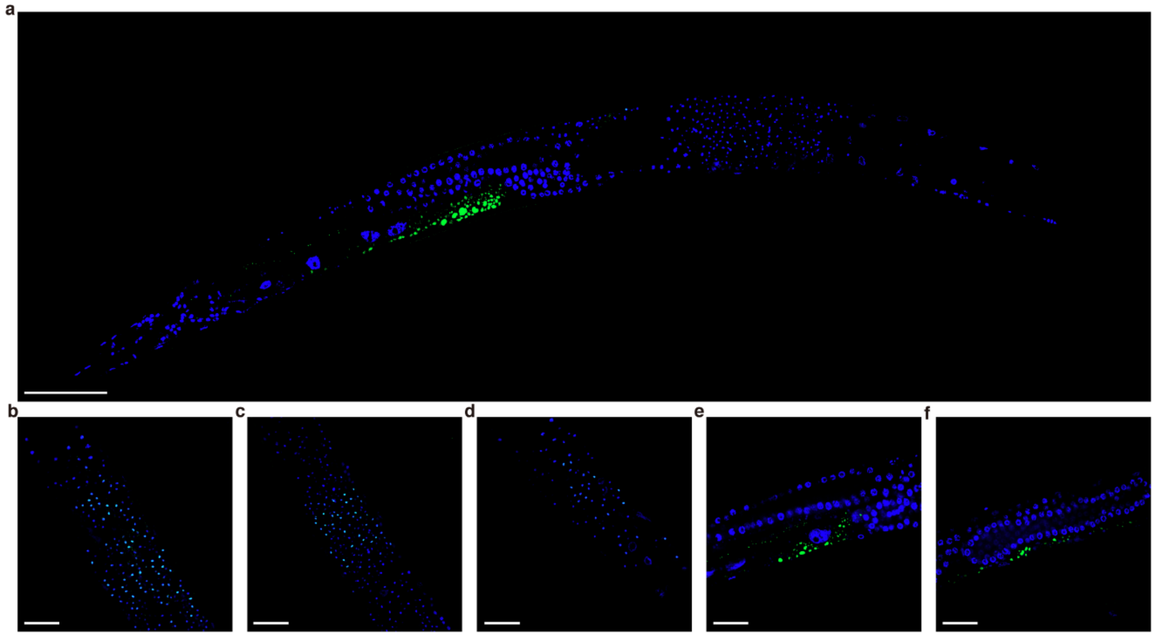


**Extended Data Fig. 8 Visualization of tissue bacteria of male worms. a**, typical image of the tissue bacteria by staining a male worm, whose tail was cut off, with DAPI and Alexa Fluor 488 dye labelled Eub338. Worms were heat-shocked at 80°C, stained at 37°C, and washed at 37°C. The excitation wavelength of the confocal fluorescent microscope was set at 405nm and 488nm, and image was obtained with a 40× oil immersion objective. Scale bar represents 50μm. **b-d**, typical images of the gonad. **e-f**, typical images of the body wall muscle. Scale bars represent 20μm.

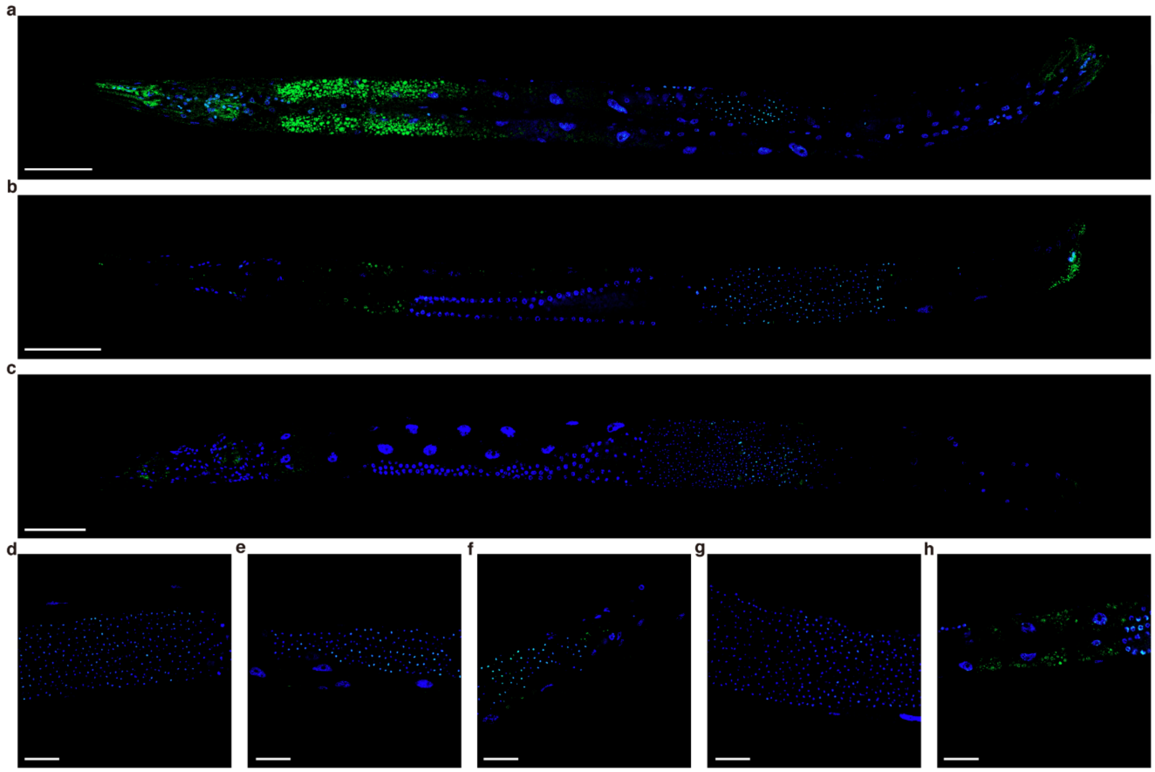

**Extended Data Fig. 9 Visualization of tissue bacteria of male worms with different FISH method. a-c**, typical images of the tissue bacteria by staining a male worm, whose tail was cut off, with DAPI and Alexa Fluor 488 dye labelled Eub338. Worm was stained at 46°C, and washed at 48°C. The excitation wavelength of the confocal fluorescent microscope was set at 405nm and 488nm. Image in **a** was obtained with a 63× oil immersion objective and **b-c** with a 40× oil immersion objective. Scale bars represent 50μm. **d-g**, typical images of the gonad. **h**, typical images of the body wall muscle. Image was obtained with a 40× oil immersion objective, and scale bars represent 20μm.

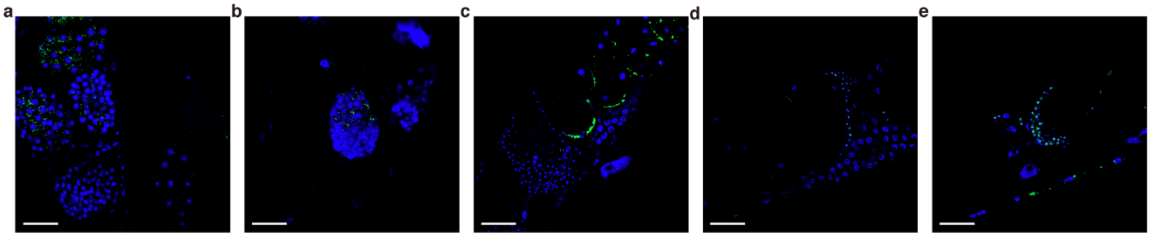


**Extended Data Fig. 10 Visualization of tissue bacteria of hermaphrodites with different FISH method. a-c**, typical images of the embryos by staining hermaphrodites, whose tail was cut off, with DAPI and Alexa Fluor 488 dye labelled Eub338. Worms were stained at 46°C, and washed at 48°C. The excitation wavelength of the confocal fluorescent microscope was set at 405nm and 488nm, and image was obtained with a 40× oil immersion objective. **a-c**, typical images of embryos in the hermaphrodites. **d-e**, typical images of the spermatheca. Scale bars represent 20μm.

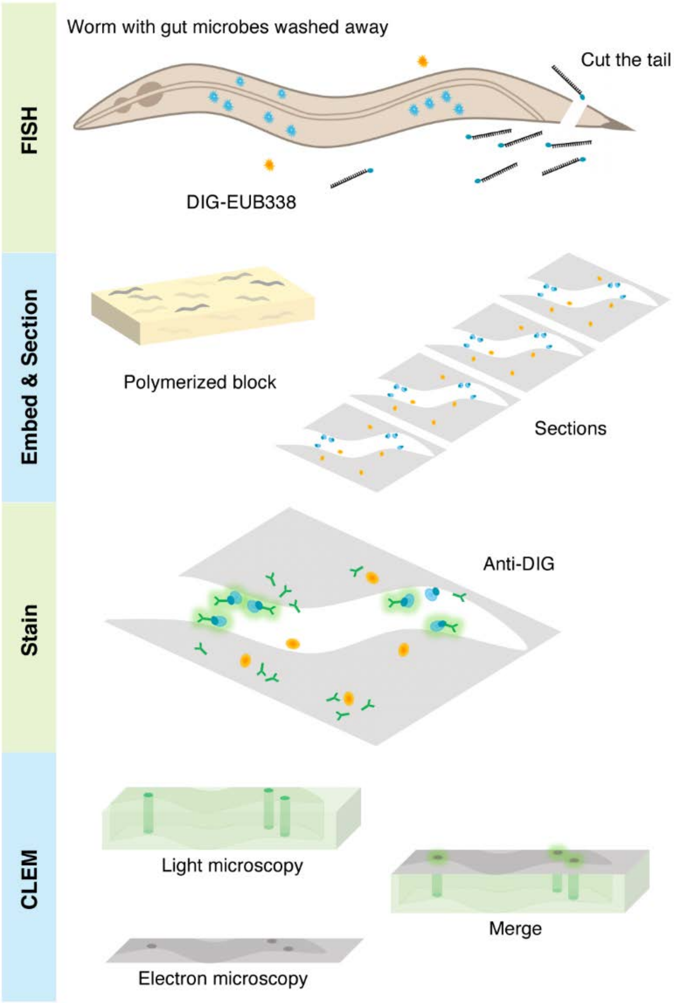


**Extended Data Fig. 11 Procedure of correlative light and electron microscopy.** In the FISH step, the worm tail was cut off and *in situ* hybridized with the digoxin labelled DNA bacterial probe. Blue indicates bacteria in the worm tissue and yellow indicates foreign bacteria. In the embedding and sectioning step, the pre-hybridized worms were washed, imbedded, and sectioned into ultrathin sections. There might be foreign bacteria (yellow) involved in this and the following steps, however, none of them was labelled with the DNA probe. In the staining step, the ultrathin section was incubated with Alexa Fluor 488 dye labelled anti-digoxin antibody (green) which specifically binds to digoxin. In the following, the sections were imaged under confocal light microscopy which exams both sides of the section, before scanned with TEM which only scans one side. Finally, these images were merged to correlate.

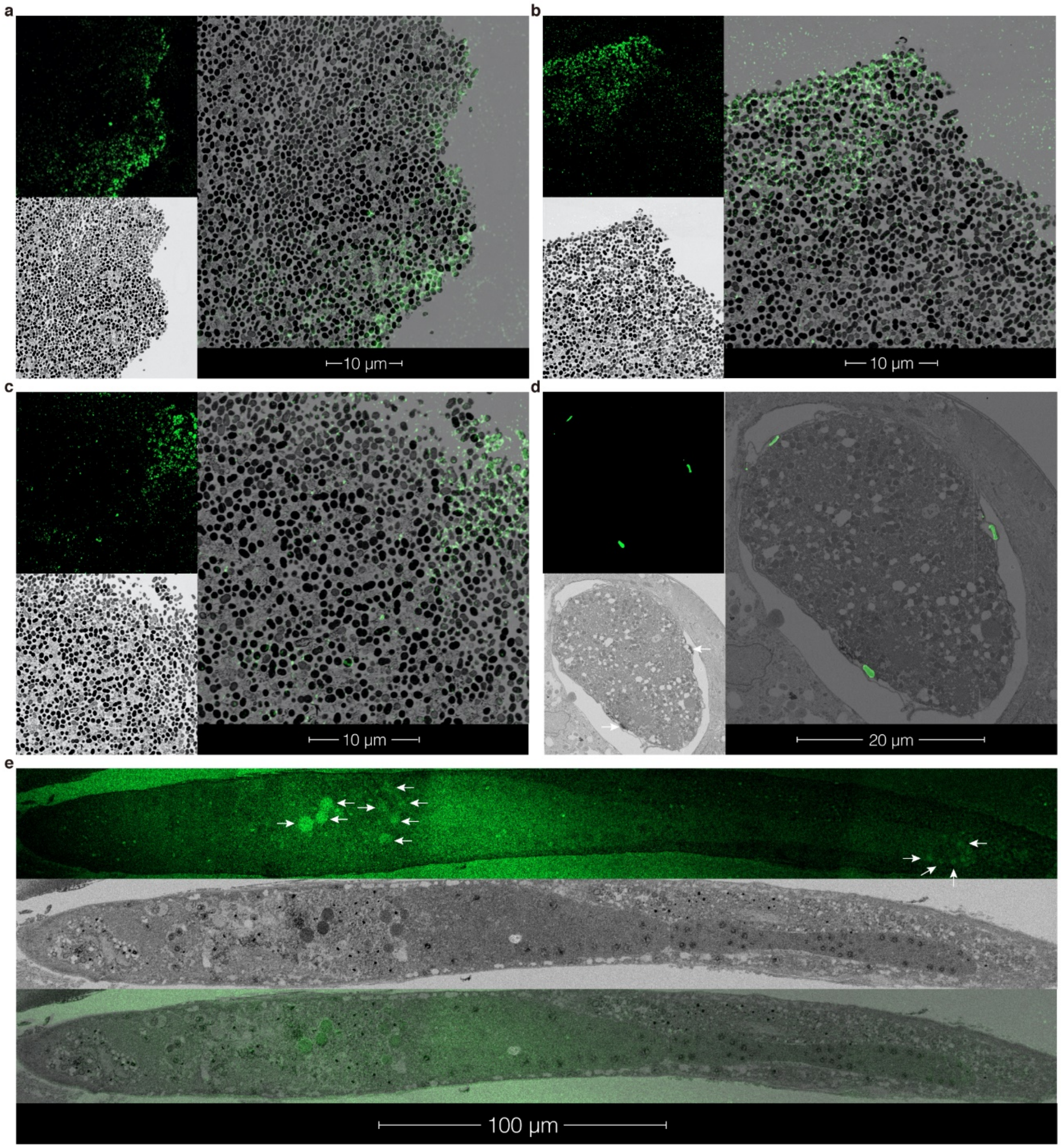


**Extended Data Fig. 12 CLEM imaging of bacteria and tissue bacteria in *C. elegans*.** **a-c**, typical images of the bacteria (*E. coli* OP50 and *E. faecalis* mix) treated with the CLEM procedure. Bacteria were stained at 46℃ and washed at 48℃. Upper left is the confocal light microscopic image of a bacteria ultrathin section obtained with a 63× oil immersion objective where the excitation wavelength was set at 488nm. Lower left is the corresponding TEM image of the same section. Right is the overlay of the LM and EM images. **d**, typical CLEM images of the embryo inside a hermaphrodite. The images are the same experiment results as in **a-c**, except the hermaphrodite was heat shocked at 80℃, stained at 37℃ and washed at 37℃. **e**, typical CLEM images of a male worm. The images are the same experiment results as in **a-c**, except the male worms were stained at 46℃ and washed at 48℃. White arrow indicates bacteria in the gonad.

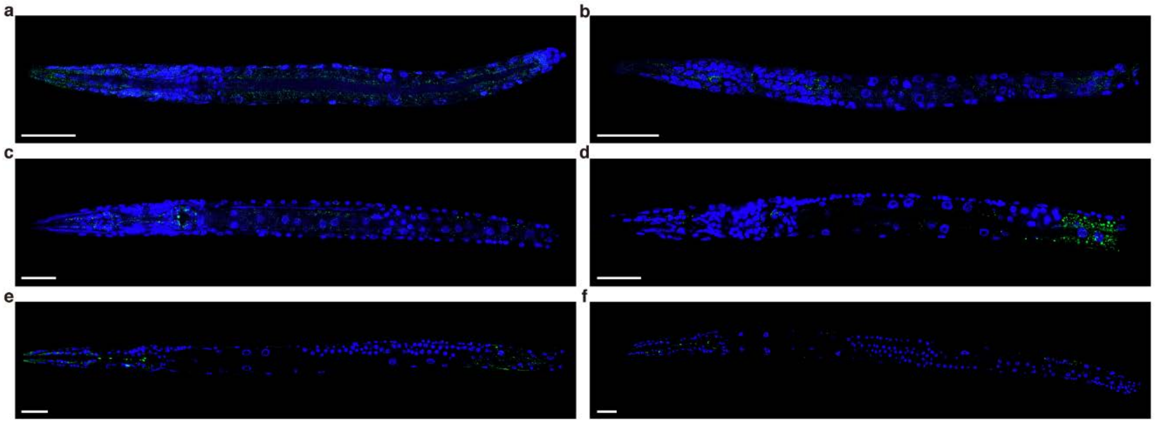


**Extended Data Fig. 13 CLEM imaging of tissue bacteria in *C. elegans* during their life cycle.** Typical images of the tissue bacteria by staining worms at different life stage, whose tail was cut off, with DAPI and Alexa Fluor 488 dye labelled Eub338. The excitation wavelength of the confocal fluorescent microscope was set at 405nm and 488nm. Worms were stained at 46℃, and washed at 48℃. **a**-**b** are images of worms at L1 stage, and **c**-**d** are images of worms at L2 stage. These images were obtained with a 63× oil immersion objective. Scale bars represent 20μm. **e** is a typical image of worms at L3 stage, and **f** is a typical image of worm at L4 stage. These images were obtained with a 40× oil immersion objective. Scale bars represent 20μm.

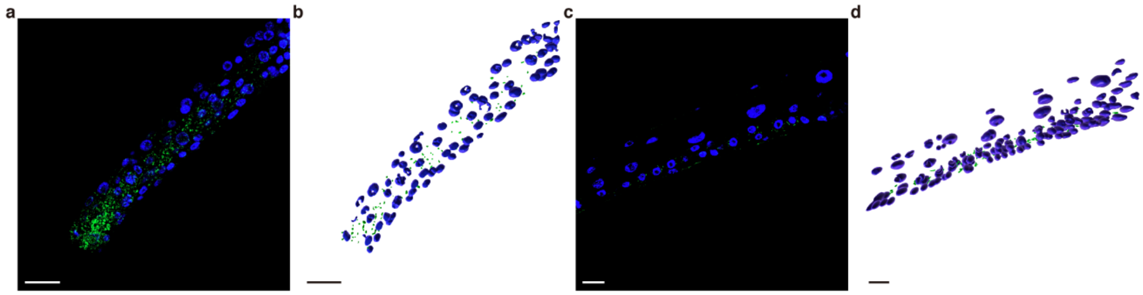


**Extended Data Fig. 14 Volumetric reconstruction of tissue bacteria in *C. elegans* during their life cycle.** Typical images of the tissue bacteria by staining worms at L1 and L4 life stages, whose tail was cut off, with DAPI and Alexa Fluor 488 dye labelled Eub338. Worms were stained at 46℃, and washed at 48℃. The excitation wavelength of the confocal fluorescent microscope was set at 405nm and 488nm. **a** is a typical image of worm at L1 stage obtained with a 63× oil immersion objective, and **c** is a typical image of worm at L4 stage obtained with a 40× oil immersion objective. **b** and **d** are the volumetric reconstructions of **a** and **c**, respectively. Scale bars represent 10μm.

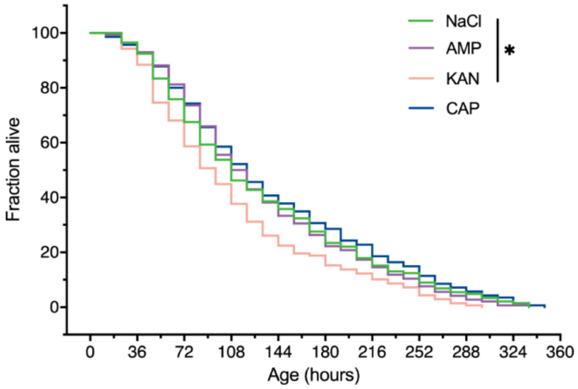


**Extended Data Fig. 15 Life span of different antibiotic treated *C. elegans* under *Enterococcus faecalis* challenging.** Life span of saline, ampicillin, kanamycin, and chloramphenicol treated worms under the challenging of pathogen *Enterococcus faecalis* ATCC29212. There are no significant differences between saline (n=145) and ampicillin (n=144) or chloramphenicol (n=140) treated worms, while kanamycin treated worms (n=138) exhibited shortened life span (Log-rank test, $P<0.05$).

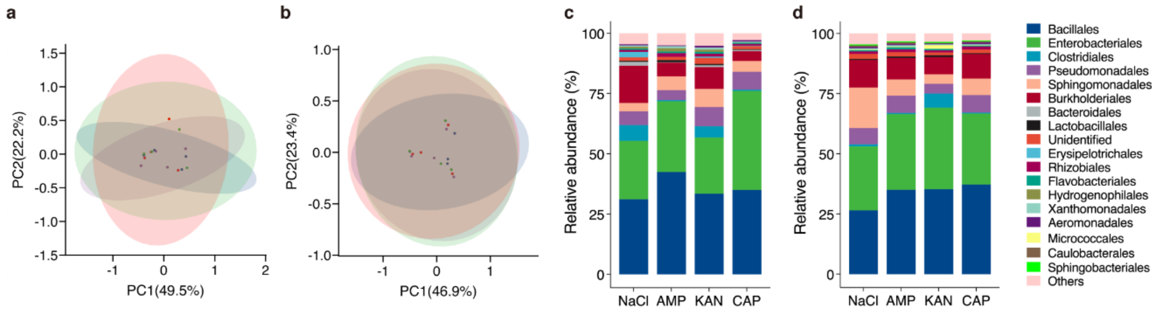


**Extended Data Fig. 16 Microbiota analysis of tissue bacteria in different antibiotic treated *C. elegans*. a,** the Bray-Curtis PCoA analysis of the tissue microbiota in the progeny of different antibiotic treated worms. **b,** the Bray-Curtis PCoA analysis of the tissue microbiota in the $2^{nd}$ generation progeny of different antibiotic treated worms. Green indicates saline treatment, purple indicates ampicillin treatment, red indicates kanamycin treatment, and blue indicates chloramphenicol treatment. For each group, n=4. **c-d**, Composition of the top 15 bacteria at the order level in the tissue microbiota of the F1 and F2 generations, respectively.

#### Supplementary Video Captions

**Video 1** *C. elegans* embryo *in situ* with bacteria inside. Blue is the DAPI stained worm embryo cell and green is the fluorescently labelled probe detected bacteria. Two embryos are shown side by side.

**Video 2** The isolated *C. elegans* embryo with bacteria inside. Blue is the DAPI stained worm embryo cell and green is the fluorescently labelled probe detected bacteria.

**Video 3** FISH detected bacteria in *C. elegans* gut. Blue is the DAPI stained *C. elegans* cell and green is the fluorescently labeled probe detected bacteria. From left to right is the head to tail of the male worm.

**Video 4** RFP labelled *E. coli* in hermaphrodite *C. elegans* gut. Red is the red fluorescent protein labelled *E. coli* ingested in a hermaphrodite worm. From top to bottom is the head to tail of a hermaphrodite worm.

**Video 5** RFP labelled *E. coli* in male *C. elegans* gut. Red is the red fluorescent protein labelled *E. coli* ingested in a male worm. From top to bottom is the head to tail of a male worm.

**Video 6** RFP labelled *E. coli* in DAPI stained *C. elegans* gut. Blue is the DAPI stained *C. elegans* cell and red is the ingested red fluorescent protein labelled *E. coli*. From left to right is the head to tail of the hermaphrodite worm.

**Video 7** *C. elegans* spermatheca with bacteria inside. Blue is the DAPI stained worm spermatheca cell and green is the fluorescently labelled probe detected bacteria.

**Video 8** *C. elegans* body wall muscle with bacteria inside at L1 stage. Blue is the DAPI stained L1 worm body wall muscle cell and green is the fluorescently labelled probe detected bacteria.

**Video 9** *C. elegans* body wall muscle with bacteria inside at L4 stage. Blue is the DAPI stained L4 worm body wall muscle cell and green is the fluorescently labelled probe detected bacteria.

#### Additional Discussion 1

##### Optimized FISH procedure

Routine FISH method not only detected bacteria in the worm gut but also in the oocyte and spermatheca, the vulva, anus, and gonad. Meanwhile, bacteria in the gut, vulva, anus and gonad could be removed by washing. This is likely because these organs all have an opening that permits the free diffusion of DNA probes. It also raised our interest to investigate whether there are bacteria in other tissues without an opening in *C. elegans*, and if so, how to detect them.

We have shown that by cutting the worm tail off, the permeability of fluorescently labelled DNA probe into the worm tissues increased so that bacteria in the tissue could be detected via FISH while foreign bacteria could not enter the tissue. Initially, the detailed FISH procedure for bacteria, the isolated embryo and worms were hybridized at 46℃ with the DNA probe and washed at 48℃ according to previous established methods[1,2]. However, we observed that bacteria in solution, the isolated embryo, and tissue bacteria in the male *C. elegans* could be visualized well with this method while tissue bacteria in the hermaphrodites could not. For hermaphrodites, the FISH procedure was finally found to hybridize with worm tissue bacteria at 80℃ and then 37℃ before being washed at 37℃ according to another procedure[3]. It is worthy to mention these temperatures were not exactly the same as in the published study since the presence of formamide in our hybridization buffer, and through this procedure the tissue microbiota could be clearly visualized in the hermaphrodites.

#### Additional Discussion 2

##### Correlative light and electron microscopy

CLEM is virtually to find the optical signal (in the light microscopy) position in the electron microscopic map of the specimen, and CLEM sample preparation is typically a compromise between optimizing light microscopy fluorescence, electron microscopy contrast and structural preservation. In our study, CLEM is performed majorly to identify whether the localization of bacteria inside or outside the worm tissue while avoiding any possible foreign contaminations.

Although we have shown that the gut and surface bacteria could be removed by extensive washing, however, it is unavoidable to have bacterial contamination in any single experimental step. In our experimental setup, the digoxin labelled DNA probe was utilized to hybridize the bacterial genes in the tail-cutoff worms in the above established FISH method. Afterwards, these labelled worms were subjected to embedding and ultra-thin sectioning. Then, the fluorescently labeled antibody against digoxin was utilized to detect the digoxin-DNA probe on the ultrathin section.

There are couple of advantages to utilize this fluorescently labeled antibody-digoxin DNA-probe coupled detection system. First, the fluorescent label was finally applied to the specimen after the embedding and ultrathin sectioning step, since the chemical process during embedding is very harsh and often damages the fluorophores. Thus, the coupled detection system avoided the chemical undermining of the fluorophores. Secondly, as we have shown in our optimized FISH method, DNA probe detects bacteria in the worm tissue without involving foreign bacterial contaminations. Thus, with the digoxin labelled DNA probe hybridized with the worm first, any bacterial contamination during the following embedding, sectioning, antibody incubation, LM observation, and EM observation does not have a chance to be labelled by digoxin attached DNA probe. Resulting that only the tissue bacteria hybridized initially before embedding could be visualized with the fluorescently labelled antibody.

However, there are some technical details that may involve discrepancies between the LM and EM observations of bacteria in worm tissues. On the one hand, it is possible that the ultrathin sectioning cut through the bacteria but did not expose its DNA. Thus, EM observation of a bacterium does not necessarily get labelled and observed under LM. On the other hand, the ultrathin section is labelled with the antibody on both sides while could only be observed under EM for one side. Thus, there could be a bacterium emitting fluorescence signal under LM, but not found under EM observations. In any case, as long as there is a fluorescence signal under LM that corresponds to a bacteria structure under EM, our demonstration the presence of bacteria in the worm tissue is accomplished.
